# Inference of Population Genetic Structure and High Linkage Disequilibrium Among *Alternaria* spp. Collected from Tomato and Potato Using Genotyping by Sequencing

**DOI:** 10.1101/827790

**Authors:** Tika B. Adhikari, Brian J. Knaus, Niklaus J. Grünwald, Dennis Halterman, Frank J. Louws

## Abstract

Genotyping by sequencing (GBS) is considered a powerful tool to discover single nucleotide polymorphisms (SNPs), which are useful to characterize closely related genomes of plant species and plant pathogens. We applied GBS to determine genome-wide variations in a panel of 187 isolates of three closely related *Alternaria* spp. that cause diseases on tomato and potato in North Carolina (NC) and Wisconsin (WI). To compare genetic variations, reads were mapped to both *A. alternata* and *A. solani* draft reference genomes and detected dramatic differences in SNPs among them. Comparison of *A. linariae* and *A. solani* populations by principal component analysis revealed the first (83.8% of variation) and second (8.0% of variation) components contained *A. linariae* from tomato in NC and *A. solani* from potato in WI, respectively, providing evidence of population structure. Genetic differentiation (Hedrick’s G’_ST_) in *A. linariae* populations from Haywood, Macon, and Madison counties in NC were little or no differentiated (G’_ST_ 0.0 - 0.2). However, *A. linariae* population from Swain county appeared to be highly differentiated (G’_ST_ > 0.8). To measure the strength of the linkage disequilibrium (LD), we also calculated the allelic association between pairs of loci. Lewontin’s *D* (measures the fraction of allelic variations) and physical distances provided evidence of linkage throughout the entire genome, consistent with the hypothesis of non-random association of alleles among loci. Our findings provide new insights into the understanding of clonal populations on a genome-wide scale and microevolutionary factors that might play an important role in population structure. Although we found limited genetic diversity, the three *Alternaria* spp. studied here are genetically distinct and each species is preferentially associated with one host.

## Introduction

The adoption of new agricultural practices and the development of new crop varieties may lead to the emergence of new pathogens or to significant changes through local adaptation in already existing pathogen populations [1, 2]. Plant pathogens continue to cause diseases in agro-ecosystems with the evolution of new races [3]. Although many plant pathogenic fungi have host preference or are host specific, some of their life histories suggest the emergence of novel pathogen species or strains adapted to new hosts [4]. Potato (*Solanum tuberosum* L.) and tomato (*Solanum lycopersicum* L.) are two important solanaceous vegetables worldwide. The center of origin of tomato and potato is believed to be South America [5,6]. *Alternaria solani* (Ell. and Mart.) Jones and Grout causes early blight (EB) on tomato and potato [7–10] whereas *A. alternata* Kessler incites leaf blight and stem canker on tomato [11–13], and brown rot on potato [14,15]. It has recently been proposed that a group of the tomato-infecting isolates was classified into a new phylogenetic species, *Alternaria linariae* (Neerg.) Simmons. This proposed split from *A. solani* (Ell. and Mart.) Jones and Grout were based mainly on multilocus sequence concatenation [16]. *A. linariae* (Neerg.) Simmons can cause EB on tomato [16–19] and potato [20].

Recent advances in high-throughput sequencing technologies offer insight into the understanding of how genome sequence variants underlie phenotype [21]. Importantly, population genomic analysis can improve our ability to develop disease diagnostic molecular tools and to resolve genetic structure, as well as better understand the molecular evolutionary history and pathogen epidemiology [22]. The use of population genomic approaches may assist in identifying genomic regions or genes that are involved in host specialization or speciation, as well as genes encoding secreted proteins that potentially interact with host molecules [23, 24]. Several secreted proteins have been identified in some model plant pathogens, and it is hypothesized that these genes rapidly evolve during host specialization due to coevolution with target host molecules [23–25].

Different molecular markers have been used to determine genetic variation in populations of *Alternaria* spp. from tomato and potato [14, 26–32]. Despite their usefulness, these markers are not appropriate for genome-wide analysis, which requires large numbers of markers. The genotyping by sequencing (GBS) approach [33], reduces the genome to fragments using restriction enzymes, and massively sequences these fragments using high throughput sequencing [33, 34]. The GBS approach directly measures loci with more than two alleles as well as short indels depending on the variant caller. This technology identifies large numbers of single nucleotide polymorphisms (SNPs) throughout the genome that are useful for genotyping and population genomic analysis [35, 36]. The GBS approach is becoming a common tool for genomics-assisted breeding in crop plants but has also been used to investigate population genetic structure, recombination, and migration in plant-pathogen populations including *Phytophthora rubi, P. infestans, P. humuli, Pseudoperonospora cubensis, Pyrenophora teres*, and *Verticillium dahliae* [34, 37, 38–41].

Genetic differentiation refers to differences in allele frequencies between populations, resulting in a significant departure from random mating in the population [42]. Evolutionary forces such as natural selection, genetic drift, and mutations or combined actions can affect genetic differentiation [43–45]. At least four measures have been used to calculate genetic differentiation between pathogen populations. Wright [42] proposed *F*-statistics (F_ST_) to measure genetic distance using biallelic markers and extended this to define genetic differences between populations. Nei [46] introduced G_ST_ for loci with multiple alleles. In addition, the *G*_ST_ is closely related to the F_ST_ and has been widely used to measure the genetic differentiation from multilocus markers such as microsatellites. With two populations and two alleles, G_ST_ ranges from 0 to 1, where 0 represents no differences in allele frequencies between two populations and 1 indicates the two populations are fixed for alternate alleles [46]. Due to the high mutation rate in microsatellite markers, *G*_ST_ can underestimate the genetic differentiation [47]. To overcome this problem, Hedrick [48] formulated a standardized measure of differentiation, called G’_ST_. This measure can standardize the observed G_ST_ value with the maximum possible value that G_ST_ could attain given the amount of observed diversity [48]. *D* measures the fraction of allelic variation among populations and can highly depend on the initial gene diversity of marker loci and the number of alleles at a locus [47]. Jost [49] introduced *D* measure and advocated that *G*_ST_ and its relatives are inappropriate measures of genetic differentiation between populations. Both G’_ST_ and Jost’s *D* become 1 at complete differentiation (even with high variation within populations) and are 0 with no differentiation [48, 49].

Linkage disequilibrium (LD) measures the non-random association of alleles among loci on the same chromosome [50]. Importantly, LD has been used to distinguish clonality from outcrossing in plant pathogens [44, 45]. Allelic associations are mostly due to physical proximity but could be affected by the mutation, recombination, natural selection, genetic drift, gene flow, and population size [51]. Several measurements have been used to estimate LD in plant pathogens. Among these, Lewontin [52] proposed deviation (*D*) as a measure of the degree of nonrandom association. Additionally, *D* compares the observed and expected allele frequencies among haplotypes and any difference between the observed and expected value is considered *D*. If two loci are in linkage equilibrium, then *D* = 0 whereas if the two loci are in LD, then *D* = −1 or +1, suggesting no evidence for recombination between the markers [45, 53]. The coefficient of correlation (r^2^) (also known as Δ^2^) is a correlation of two alleles at the two sites [54]. The r^2^ or Δ^2^ values range between 0 and 1, where 1 represents when the two markers provide identical information and 0 when they are in perfect equilibrium [54]. Hedrick [53] compared coefficients of LD and found that *D*’ estimate was better than other estimates. Another measure of LD at a multilocus scale is the index of association (I_A_) [55] and a standardized multilocus index of association (*ř*_d_), which is an alternative measure of I_A_ [56]. Three balancing factors such as genetic drift (a function of population size), random mating, and distance between markers limit linkage equilibrium or LD [45, 53, 54]. The genetic diversity of *A. alternata* and *A. solani* from tomato and potato has been reported from Brazil [28, 57], China [59], Cuba [29], and Wisconsin [14, 31]. However, little is known about the population structure of *A. linariae*, although microsatellite loci and gene genealogy approaches have been used recently [26, 27]. We hypothesize that *A. linariae* is more prevalent and endemic to high altitude and cold environment in tomato production areas in western in NC. Thus, one of the main goals of the present study was to scan genome-wide diversity and compare populations of *A. linariae* collected from tomato in NC with *A. solani* from potato in WI.

In a previous study, population genetics of *Alternaria* spp. collected from tomato and potato was examined using microsatellite loci. Our results revealed high levels of gene and genotypic diversity and mixed reproduction (both asexual and sexual) in the populations analyzed [26]. In another study, a coalescent gene genealogy analysis revealed no evidence of sexual recombination in *Alternaria* spp. [27]. To resolve these issues, we tested two hypotheses in this study. First, we tested the hypothesis that populations of *Alternaria* spp. are genetically distinct, in order to support the previous observations that these species are differentiated by host preference [28]. Second, we used LD analysis to reassess the hypothesis that *Alternaria* spp. are predominantly clonal with no evidence of sexual recombination. Here we address the following questions: (i) how polymorphic are natural populations of *Alternaria* spp., as revealed by GBS? (ii) what degree of genetic differentiation can be partitioned between the two morphologically similar sister species: *A. linariae* and *A. solani*? (iii) if present, what is the extent and degree of LD? To answer these questions, we genotyped and sequenced a panel of 187 isolates from three *Alternaria* spp. collected from tomato and potato from NC and WI using GBS.

## Materials and Methods

### Ethics statement

This research was conducted in accordance with the guidelines and recommendations of the United States Department of Agriculture Animal and Plant Health Inspection Service permit P526P-14-00008.

### The species and populations

*Alternaria* spp. are haploid and molecular evidence indicates that they undergo sexual reproduction although a teleomorph has never been found [60]. The isolates of *Alternaria* spp. (*A. alternata*, *A. linariae*, and *A. solani*) from naturally infected tomato and potato fields in NC and WI were chosen from our collections and were described in previous publications [26, 31]. A population herein is defined as a geographic location where the isolates of *Alternaria* spp. were collected [34]. The populations collected from these locations were adapted to local climatic conditions (temperature and elevation), hosts (tomato and potato) and genetic stocks (breeding lines, hybrids, heirloom), agricultural practices and length of growing seasons (Table 1). A panel of 187 isolates of *Alternaria* spp. was selected and analyzed by GBS in this study (Supplementary Table 1). Among them, *A. alternata* isolates (*N* = 42) were mainly collected from heirloom tomatoes from central, eastern, and southwestern NC. Populations of *A. linariae* consisted of 95 isolates collected from fresh market hybrid tomatoes from four counties in western NC. These populations composed of Haywood county (*N* = 31), Macon county (*N* = 23), Madison county (*N* = 20) and Swain county (*N* = 21). *A. solani* isolates (*N* = 50) were collected from potato fields from Waushara county in WI.

### Genomic DNA extraction and preparation

For each experiment, mycelial plugs of the isolates were revived from −80°C by plating on acidic potato dextrose agar (A-PDA, 39 g/L, 50 mg/L streptomycin). To curtail systematic differences in the treatment of isolate, the culture medium was prepared the same day and handled by the same person for each of the experimental steps as described below. A single conidium of each isolate was transferred to new A-PDA plates. Cultures were grown at 26°C for 1 week and harvested. Hundred mg of mycelium tissue was lyophilized and ground to a fine powder in 2-mL tubes with glass beads using a Tissue Harmonizer (Model D1030-E, Beadbug™, Benchmark Scientific Inc., Edison, NJ, U.S.A) at 400 rpm for 2 min. Genomic DNA of each isolate was extracted using the DNeasy^®^ Plant Mini kit (Qiagen, Germantown, MD, U.S.A.) following the manufacturer’s instructions. The DNA concentration was quantified using a NanoDrop spectrophotometer ND-1000 (NanoDrop Technologies Inc., Wilmington DE, U.S.A). Approximately 20 ng of each DNA sample was digested with the restriction enzyme Eco*RI* (New England Biolabs, Ipswich, MA, U. S. A.) at 37°C for 2 hours and quality was evaluated on 1%agarose by gel electrophoresis. After checking the quality, 100 ng/μL of purified DNA of each isolate was transferred into 96 well-skirted plates (95 samples and one blank well). All 187 DNA samples were split into two plates and shipped to Cornell University, Institute for Genomic Diversity (IGD) for GBS analysis (http://www.igd.cornell.edu/index.cfm/page/GBS.htm).

### Normalization and selection of restriction enzymes (REs)

To select a suitable RE to recognize the *Alternaria* DNA sequence, libraries for GBS were generated according to the protocol described previously [33]. Genomic DNA of *A. solani* isolate # BMP 0185 from which the draft reference genome generated [61]; http://alternaria.vbi.vt.edu), was kindly provided by Prof. Barry M. Pryor, University of Arizona, U. S. A. Genomic libraries were generated using two REs: *Ape*KI (5-bp recognition site) and *Pst*I (6-bp recognition site). *Ape*KI yielded more genomic coverage than *Pst*I (*data not shown*) and was selected to identify SNP variants in genomic regions of *Alternaria* spp.

### Genotyping by sequencing (GBS)

Library preparation and sequencing were performed at Cornell University, IGD as described previously [33]. Briefly, each DNA sample was digested with *Ape*KI, and adapters were ligated to each DNA sample. The samples were pooled, PCR-enriched, and purified before being sequenced with 100-bp single-end using the Illumina Hi-Seq 2500 sequencing platform (Illumina, San Diego, CA, U.S.A.).

### Read mapping and SNP calling

The variant calling files (VCF) resulting from Illumina sequencing were obtained from IGD and processed as described previously [33]. Reads were mapped to both draft reference genome sequences of *A. solani* isolate # BMP0185 and *A. alternata* isolate # BMP0270 [61]; http://alternaria.vbi.vt.edu) and were further trimmed to remove low quality. Raw sequencing reads were processed with the TASSSEL-GBS analysis using a TASSEL Version 3.0.173 pipeline [62]. Variants resulting from the TASSEL pipeline were treated as raw variants and were received as a variant call format (VCF) file [63].

### Variant data analysis

Due to the evolutionary divergence among the sampled taxa, and our ability to call homologous loci, the SNP variants were divided into three data subsets. First, the raw variants that resulted from the TASSEL pipeline were used to compare *A. alternata* with *A. linariae* and *A. solani*. Second, the TASSEL variants that resulted from mapping reads to the *A. solani* genome were processed by calculating 10^th^ and 90^th^ percentiles of sequence depth at variable positions for each sample, marking genotypes above or below these percentiles as absent, as well as any variants that had a depth less than 4. The TASSEL pipeline produced diploid genotypes for this organism that is expected to be haploid. To create a haploid data set, any heterozygous positions were deleted from the data set while homozygous positions were scored as one allele. These variants were then filtered to include only variants with less than 5% missing data. Third, the quality-filtered data from the second data set was subset to only the samples from four counties in NC and then the variants were subset to only include variable sites. These processing steps were performed in R [64] using vcfR [65]. To compare *A. alternata* to *A. linariae* and *A. solani*, the number of missing genotypes were used. The individual percent of missing genotypes were calculated from the variants that were mapped to the *A. alternata* draft genome and *A. solani* draft genome. This data was summarized in a bar-plot created in R [64]. To discard variants due to errors, we created quantiles of sequence depth for each sample. We omitted SNP variants that were in the lowest 10% of coverage and the highest 10% of coverage.

### Principal component analysis (PCA)

Patterns in genome-wide relatedness among and within species of *Alternaria* were performed by PCA [66]. To compare *A. linariae* to *A. solani* populations, the VCF data were converted to a genlight object using vcfR::vcf2genlight() and analysis was performed with adegenet::glPca() [67, 68]. Results from this analysis were plotted using ggplot2 [69]. To compare among populations of *A. linariae* collected from a tomato from four counties in NC, we also used PCA as reported above.

### Genetic differentiation

We used Hedrick G’_ST_ [48] to quantify the degree of genetic differentiation among populations of *A. linariae* from four counties in western NC. Genetic differentiation was estimated by calculating G_ST_ [46, 70] and G’_ST_ [48] using vcfR::pairwise_genetic_diff() (Knaus and Grünwald 2018) [71]. To visualize linkage throughout the genome, Manhattan plots were created using vcfR [65].

### Clonality and linkage disequilibrium (LD) estimation

For LD estimation, isolates from each region or county were initially considered as one population sample. LD is the non-random association of alleles at adjacent loci and was calculated using Lewontin’s *D* [52]. To infer the mode of reproduction (e. g., clonal or sexual), we used VCF tools to calculate Lewontin *D* and squared it so it would range from 0 to 1. To quantify the LD decay, we plotted Lewontin *D* against physical distance using R. We also used an LD heatmap (http://stat.sfu.ca/statgen/research/ldheatmap.html) to visualize linkage on exemplar contigs and also calculated associations between alleles at different loci.

## Results

### Comparison among *A. alternata*, *A. linariae*, and *A. solani* populations

The raw VCF files produced by TASSEL consisted of 152,114 variants. Of the 187 isolates of *Alternaria* spp. genotyped, three isolates: P14, T20, and To3 were found to be of low quality based on the degree of missing genotypes reported by TASSEL. Therefore, these three isolates were not included in further analyses. Approximately the same size, 31-33 Mbp (*data not shown*) were detected when the isolates of *Alternaria* spp. sequences were compared with both draft reference genome sequences of *A. solani* isolate # BMP0185 and *A. alternata* isolate # BMP0270 [61]. These genomes appeared to have equal representation of GC and AT content and both have a peak of GC content (*data not shown*). To compare three *Alternaria s*pp. at the genome level, reads derived from TASSEL were mapped to the *A. solani* BMP0270 reference genome and had 10 to 20% missing variants or SNPs regardless of the host (tomato or potato). However, when reads that were mapped to the *A. alternata* reference genome, the *A. linariae* and *A. solani* isolates contained 50 to 60% of missing variants (Fig. 1).

**Figure 1.**
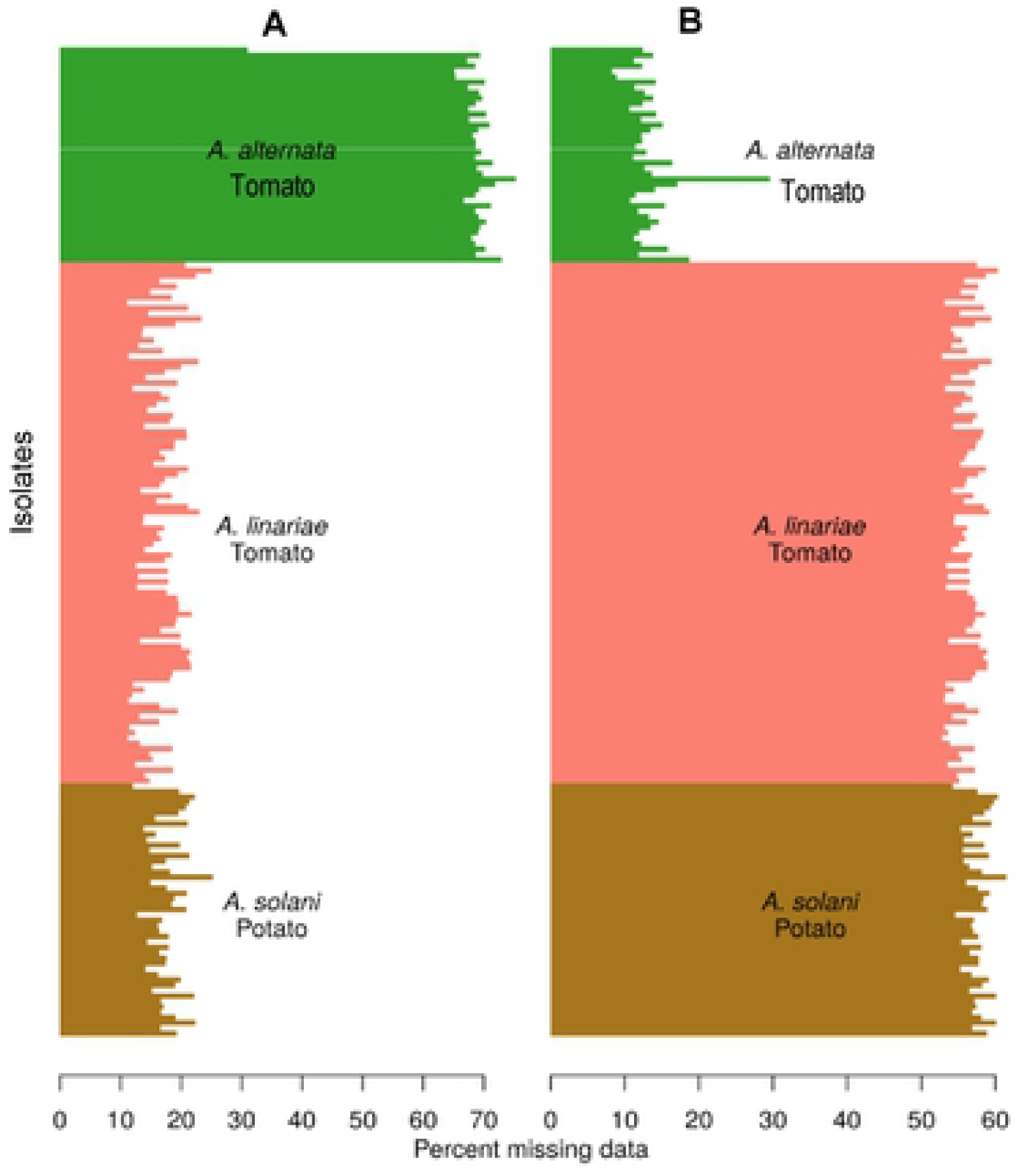
Genome-wide comparisons *A. alternata* versus *A. linariae* and *A. solani*. Reads derived from TASSEL were compared and aligned to the *A. alternata* draft reference genome strain BMP0270 (**A**) and the *A. solani* draft reference genome strain BMP0185 (**B**) [61]; http://alternaria.vbi.vt.edu). Data from isolates of *A. solani* had low (10 to 20%) missing variants regardless of the host (tomato or potato). However, when *A. alternata* mapped to the *A. solani* reference genome resulted in a high (60 to 70%) missing variants.

### Principal component analysis (PCA)

#### Comparison between *A. linariae* and *A. solani*

We tested the hypothesis that morphologically similar two *Alternaria* spp: *A. linariae* and *A. solani* are genetically distinct. Ordination of the isolates using PCA demonstrated that the *A. linariae* and *A. solani* taxa were well differentiated. The first and second components accounted for 83.9% and 8.0% (total of 91.9%) of the variation explained (Fig. 2). The first axis differentiated among isolates of *A. linariae* collected from tomato while the second axis consisted of *A. solani* collected from potato.

**Figure 2.**
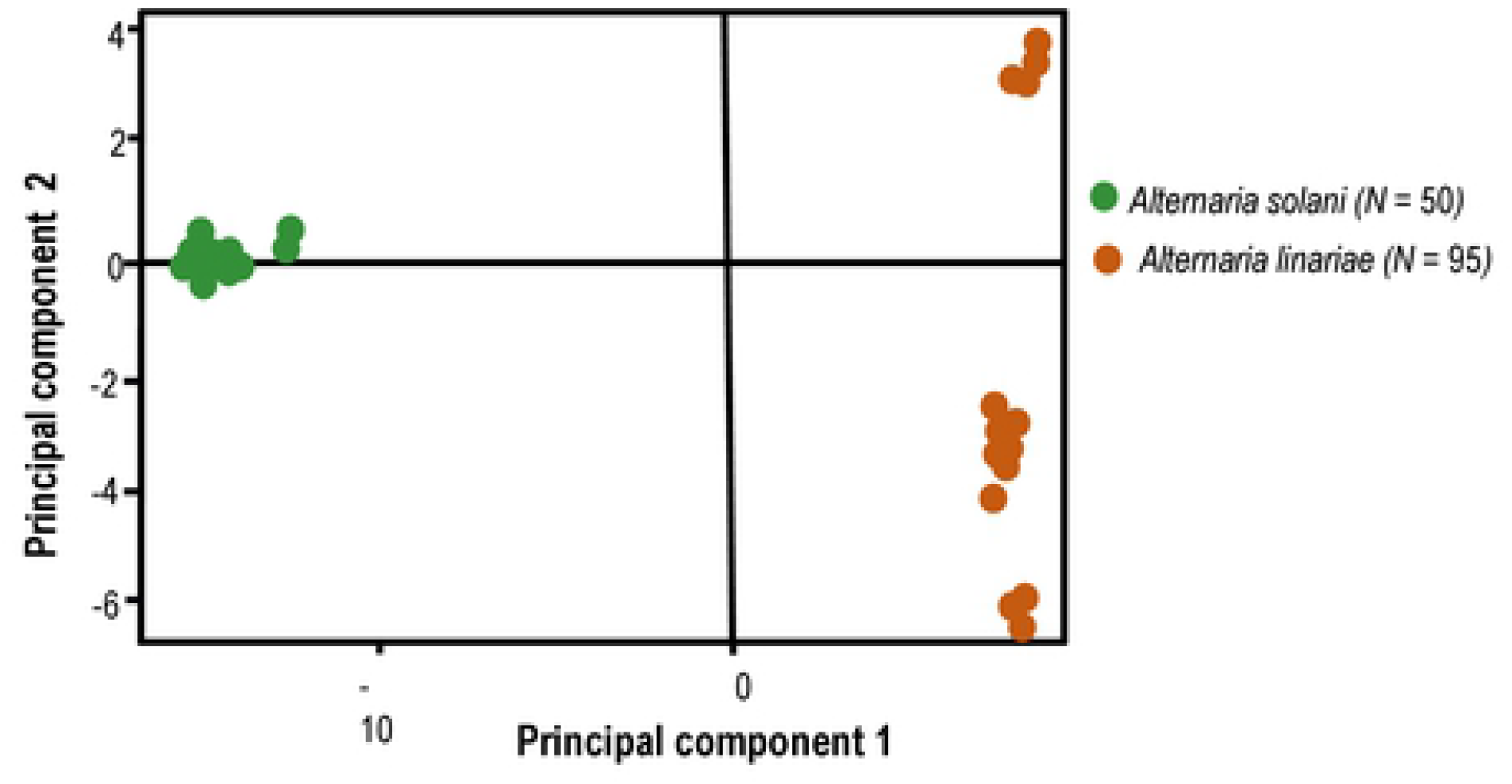
Principal components analysis of isolates of *Alternaria solani* collected from potato in Wisconsin and isolates of *Alternaria linariae* from tomato. Populations were highly differentiated by pathogen by the first component. Axes one and two accounted for 83.9% and 8.0% of the variation in the data, respectively.

#### Comparisons among *A. linariae* isolates

We tested the hypothesis of genetic differentiation across populations *A. linariae* in NC. PCA revealed three clusters of *A. linariae* isolates; axes 1 and 2 of the PCA accounted for 69.30% and 28.70% of the total genetic variation (Fig. 3). PCA also indicated that the populations of *A. linariae* from four counties in NC were clustered in three groups. Comparisons among four populations of *A. linariae* from NC analyzed for genetic differentiation by genomic position ranged from no differentiation to highly differentiated. Isolates of *A. linariae* collected from Haywood and Macon counties showed no differentiation (G’_ST_ = 0.0) (Fig. 4A). Variants obtained for isolates collected from Haywood and Madison counties appear to fall into two classes (Fig. 4B). Many variants showed no differentiation (G’_ST_ = 0.0). The second class of variants exhibited moderate differentiation (G’_ST_ = 0.15). Similarly, isolates from Macon and Madison counties had variants that fell into two classes (Fig. 4C). Some variants demonstrated no differentiation (G’_ST_ = 0.0) while others were well-differentiated (G’_ST_ = 0.2). Isolates collected from Haywood and Swain counties had variants that fell into three classes (Fig. 4D). Some variants were completely differentiated (G’_ST_ = 1.0), highly differentiated (G’_ST_ = 0.9), or moderately differentiated (G’_ST_ = 0.15). Isolates collected from Macon and Swain counties had variants that fell into three classes (Fig. 4E) and all of them were differentiated (G’_ST_ = 0.2 to 1.0). A comparison of isolates collected from Madison and Swain counties had variants that demonstrated nearly complete differentiation (G’_ST_ > 0.8) (Fig. 3F).

**Figure 3.**
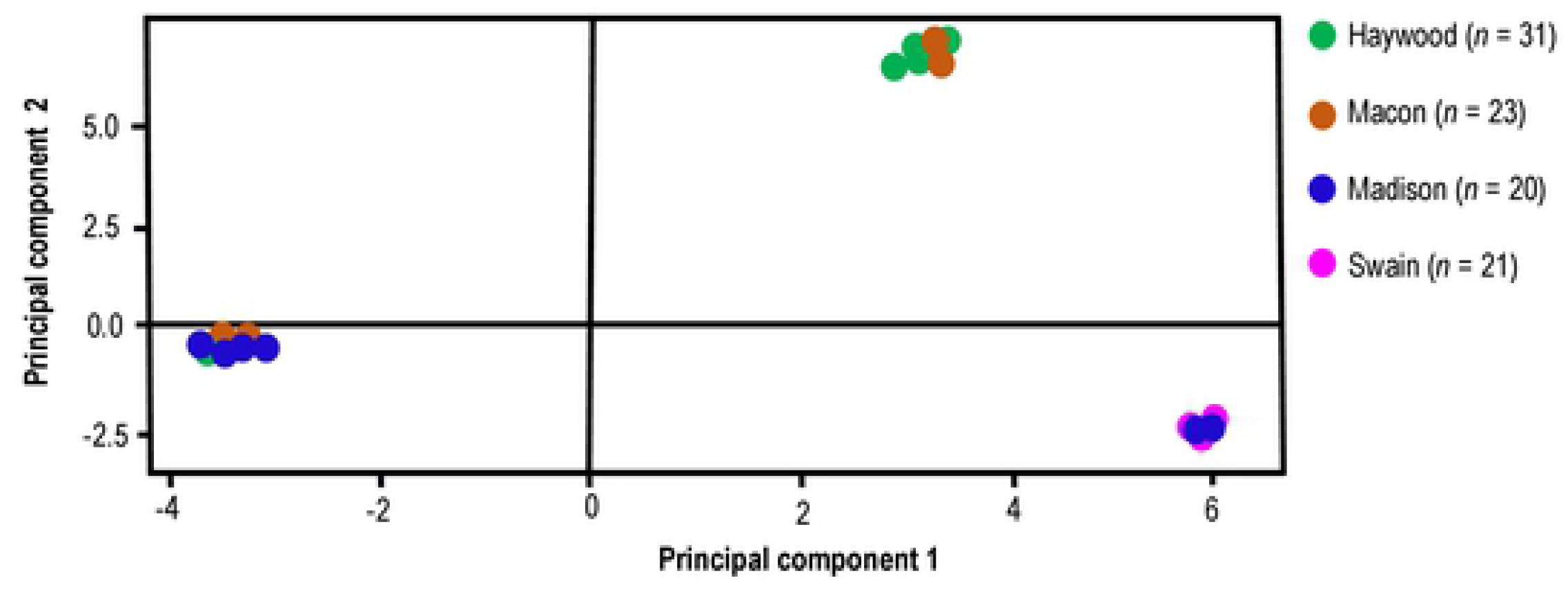
Principal components analysis of *Alternaria linariae* populations collected from tomato in four contiguous counties: Haywood, Macon, Madison, and Swain in North Carolina. Shown are three genetic groups. Axes one and two accounted for 69.30% and 28.70% of the variation in the data set, respectively. The isolates in the right quadrants at the top of the first principal coordinate were from Haywood and Macon counties and the isolates at the bottom were from Macon and Swain counties. Similarly, the left quadrants consisted of the isolates from Haywood, Macon, and Madison counties (Fig. 3).

**Figure 4.**
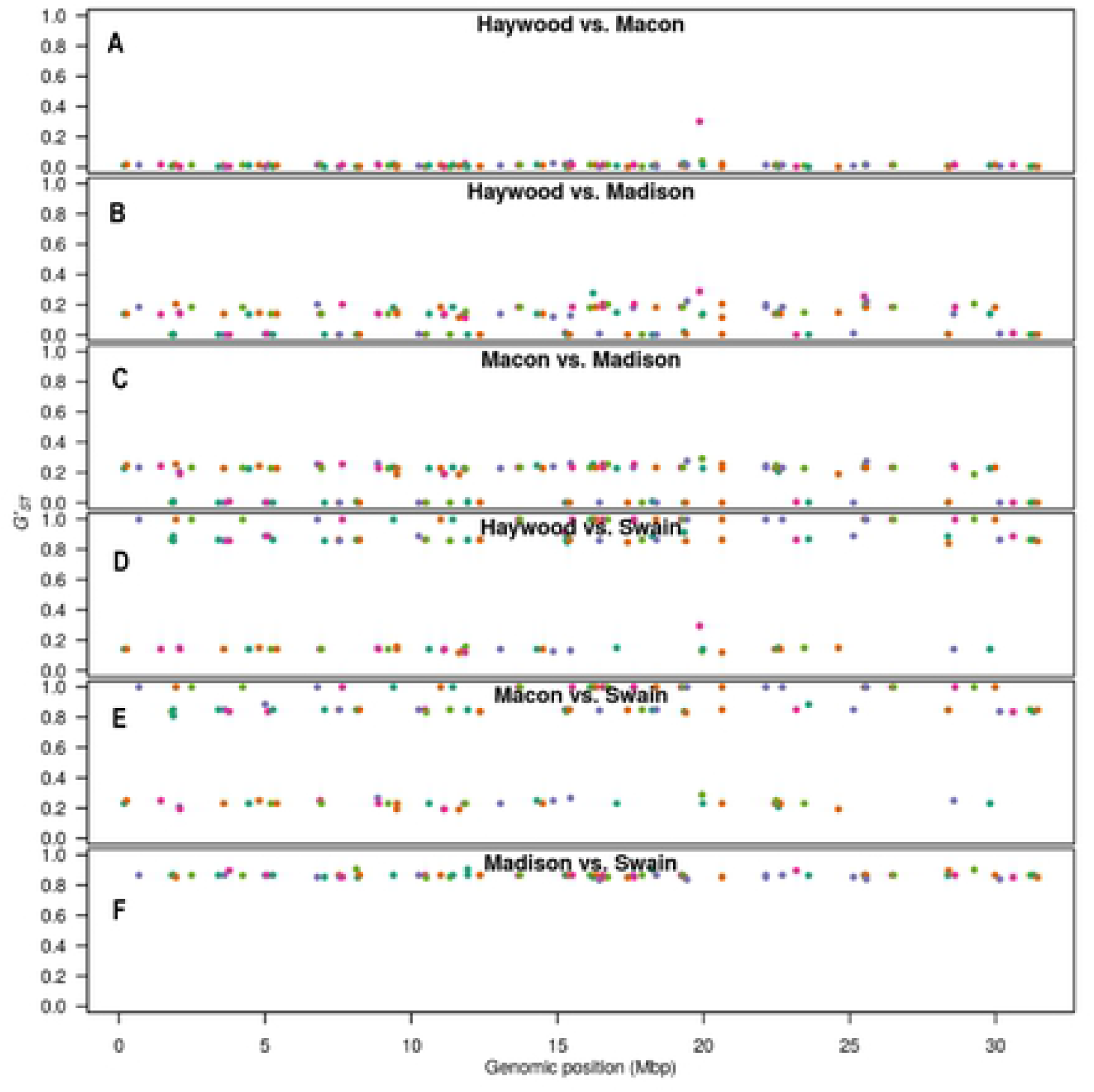
Manhattan plot showing little or no differentiation among populations of *Alternaria linariae* in North Carolina. Each variant was color-coded according to the contig throughout the genome and it came from using a recycled eight-color palette. Contigs were ordered sequentially from largest to smallest. Hedrick’s G’_ST_ [48] estimate of population differentiation is presented on the y-axis while the cumulative genomic position is presented on the x-axis. Genetic differentiation is shown pairwise among populations of *A. linariae* collected from four counties: Haywood, Macon, Madison, and Swain in North Carolina. Isolates from Swain county were highly differentiated (G’_ST_ > 0.8) throughout the genome whereas isolates from Haywood, Madison, or Macon counties are undifferentiated or little differentiated (G’_ST_ 0.0 - 0.2).

### Inference of linkage disequilibrium (LD)

To infer the mode of reproduction (e. g., clonal or sexual) in *Alternaria* spp., we compared contigs with the greatest number of variants. In general, LD showed no apparent relationships. Pairwise comparisons between Lewontin *D* [52] values and physical distances suggest a lack of recombination in the *Alternaria* spp. (Fig. 5A). Instead, we observed very high levels of linkage regardless of the physical distance, possibly due to asexual reproduction. We used G’_ST_ to find highly differentiated SNP variants and visualized these. To further confirm the population differentiation due to novel alleles, we estimated the allelic composition of each population. The reference allele, 0 and the first alternate allele, 1 appear subjectively similar in each population (Fig. 6). However, the second alternate allele, 2 in the Swain population (0.0245) is 2.5 times more abundant than it appears in the population with its second greatest abundance (Macon population: 0.0096).

**Figure 5.**
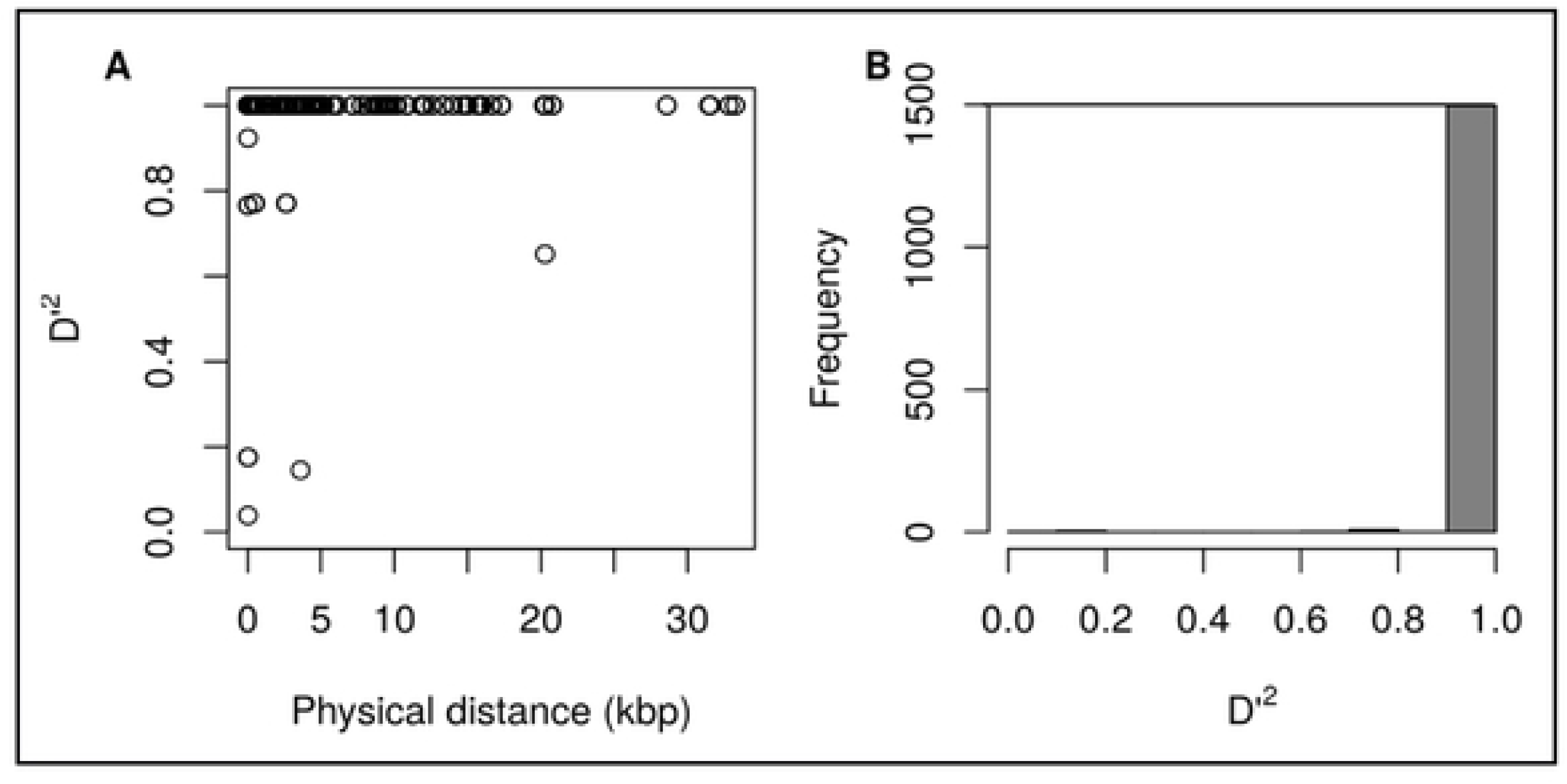
Visualization for linkage disequilibrium (LD) measured as Lewontin *D* [52]. Linkage does not decay with increasing physical distance. Loci that are close together are expected to have relatively high linkage and this is expected to decay with increasing physical distance due to recombination events. The lack of LD decay is indicative of isolates that are reproducing asexually or clonally. Lewontin [52] *D* values of 0 indicate linkage equilibrium where values of 1 indicate LD.

**Figure 6.**
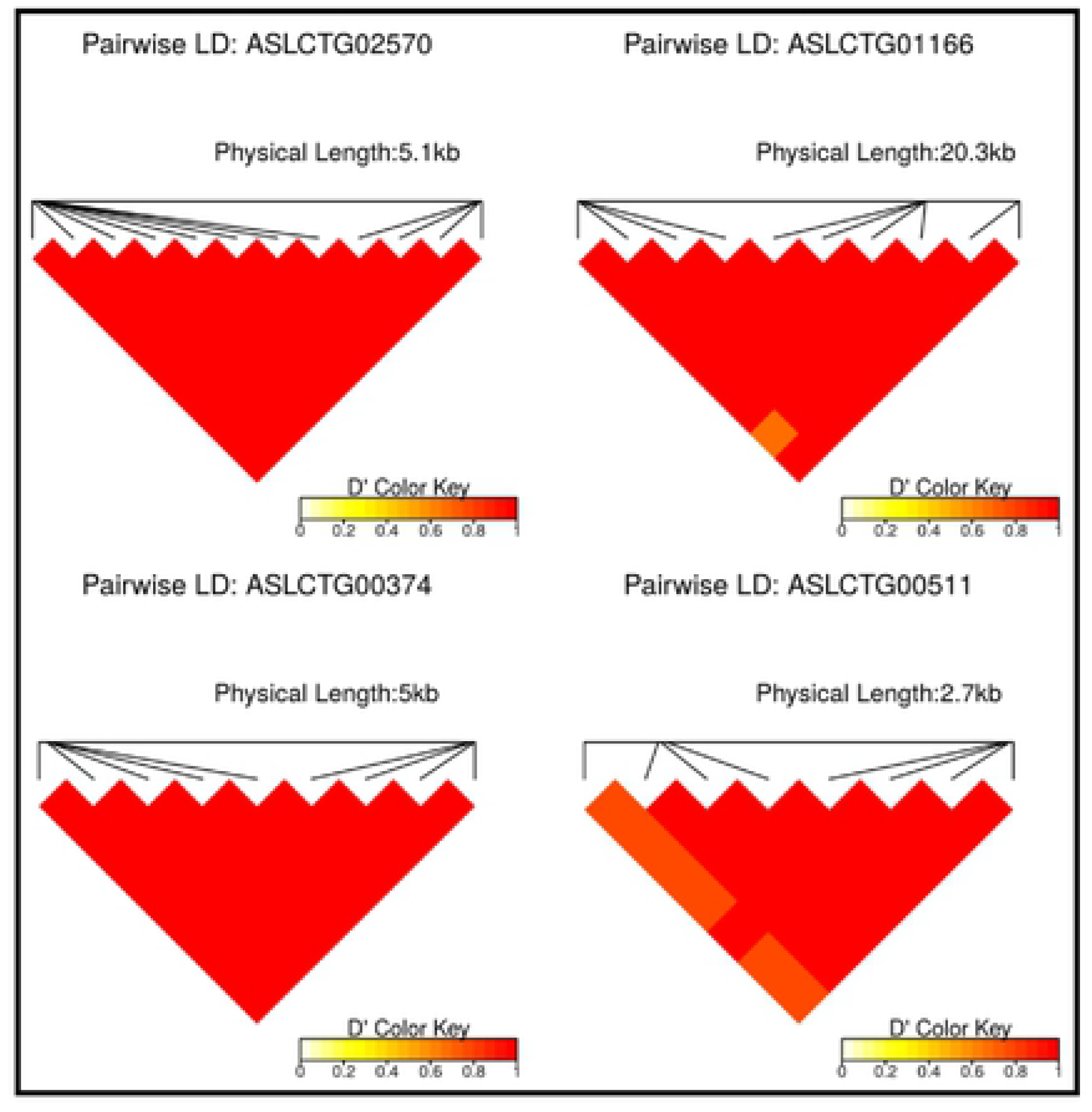
Pairwise comparisons of variants with high G_ST_ values in the four longest contigs with the highest number of variable sites. Linkage disequilibrium occurs throughout each entire contig. The contigs with the greatest number of variants demonstrated almost complete linkage throughout their length.

## Discussion

We investigated the potential utility of the GBS approach for the determination of allele frequencies in three closely related *Alternaria* spp. collected from NC and WI. Since three species of *Alternaria* cause diseases on both tomato and potato, we were interested in mapping SNPs in their genomic regions. In doing so, we mapped our test isolates to both *A. alternata* and *A. solani* draft reference genomes [61]. We observed a dramatic difference in genotype calling performance, resulting in two sets of SNP variants. When the isolates of *A. alternata* were mapped to the *A. solani* draft reference genome, a low number of *A. alternata* isolates passed quality control and consisted of a high percentage of missing SNP variants. Thus, a comparison of *A. alternata* with *A. linariae* and *A. solani* appeared to be challenging. As a result, our analysis was focused on two species: *A. linariae* and *A. solani*. Data analysis further demonstrates that variant combinations observed in natural populations of *Alternaria* spp. are not random samples. SNP variants act as one locus in the genome and our results provide evidence of an excess of homozygosity within the populations that can reproduce asexually [72]. Our results also indicate that genetic differentiation values are fixed throughout the genomes of the isolates analyzed and these results support the hypothesis that there was little or no genetic differentiation within *A. linariae* populations collected from NC.

Historically, host preference and morphological features have been used as principal criteria for species delimitation in *Alternaria* spp. [8, 10]. More recently, the gene genealogy approach was used to reclassify species of *Alternaria* [16]. Particularly, two species: *A. linariae* and *A. solani* can infect either tomato or potato, or both [9, 16, 20]. To examine specific population structure and the genetic differentiation, we represented the isolates of *A. solani* and *A. linariae* by host origins. Although these two fungal pathogens coexist in the same agro-climatic conditions and cropping systems, the disease management strategies, resistance breeding approaches, and evolutionary history are different or species-specific [8, 26, 73].

Our data indicate differentiation of the isolates of *A. solani* collected from a potato from WI and the isolates of *A. linariae* collected from a tomato from NC could be due to geographic isolation, which is ~1560 km apart from each other. Doyle et al. [74] implemented an intensive sampling strategy and collected *Colletotrichum gloeosporiodes* sensu lato (*s. l*.) species complex of plant pathogens and endophytic fungi from wild and cultivated cranberry production regions in North America. They found that the hosts’ habitats and the hosts’ species from which an isolate has been isolated were the useful indicators of species identity in the sampled region. In contrast, Meng et al. [59] collected four populations of *A. solani* from diverse potato production regions in China and analyzed with microsatellite markers. Although distinct clones were detected in the populations which were separated by thousands of kilometers, the random association among SSR loci was detected in half of the populations assessed. Intriguingly, there was no separation of *A. solani* populations in China by distance [59]. Our study showed that host, surrounding habitat of the host, edaphic factors, and nature of the fungus-host association could be predictive for the population structure of *A. solani* and *A. linariae*.

We further quantified genome-wide genetic differentiation among populations of *A. linariae* collected from tomato in four contiguous counties in western NC. Apparently, low genetic diversity was observed in all populations of *A. linariae*. The Swain County population appeared to be the population most differentiated from the others and it had the lowest genetic diversity. Although repeated samplings would make direct comparisons with populations of *A. linariae*, from different counties in NC, none of the samples from four contiguous counties in NC formed a single cluster in PCA analysis. This result indicated that the isolates of *A. linariae* either originated from the same ancestral population without additional genetic diversity resulting in fixed alleles or by a regular admixture of the clades differentiated in the PCA [75]. A similar finding was reported for *Sclerotinia sclerotiorum* populations collected from *Ranuculus ficaria* (meadow buttercup) in Norway which were described by low genetic diversity [75, 76].

One of the significant findings of the present study is that there was no relationship between LD decay and physical distance. We found a lack of pairwise LD decay with the increasing physical distance between variant loci. The contigs with the greatest number of variants demonstrated almost complete linkage throughout their length. When the entire genome is linked, due to asexuality, selection cannot act on individual genes but instead acts on the entire genome as if it were one locus. Our previous study based on coalescent genealogical approaches also revealed a lack of recombination breakpoints in *Alternaria* spp., suggesting strongly clonal populations in NC and WI [27].

A variety of evolutionary processes such as epistatic selection, hitch-hiking, admixture, and physical linkage among markers can cause LD [45, 78]. For example, epispastic selection has been reported as the main factor responsible for generating LD in *Blumeria graminis* f. sp. *hordei* (barley powdery mildew pathogen) where the use of selective fungicides and resistant cultivars favored the selection of avirulence or fungicide resistant alleles during the asexual stage [79, 80]. Future research should be directed to select few isolates from dominant haplotypes of *A. linariae* and address whether these isolates have greater fitness attributes to adapt in environmental conditions and fungicide resistance and have high levels of reproductive capabilities and dispersal abilities.

Another possible cause of LD can be population admixture, which is sampling genetically different populations and analyzing them as one population [78]. Using microsatellite loci, we found high genotypic diversity, multilocus genotypes, and random association alleles in a few populations of *Alternaria* spp. collected from tomato and potato from NC and WI, respectively (Adhikari et al. 2019a). Although there was no evidence of an active sexual cycle in *Alternaria* [60], both mating-type idiomorphs have been detected in some populations of *Alternaria* spp. analyzed [18, 26, 58, 77]. Our previous study [27] also indicated that the clonal population structure of *Alternaria* spp. resulted from a combination of asexual reproduction predominantly via conidia and haploid selfing of sexual recombination [75].

Genetic drift alone can create LD between closely linked loci (Slatkin 2008). Importantly, genetic drift is a [81]. Moreover, the effects of drift are therefore strongest in small populations, in which a few events can have a large impact. We further hypothesize that potential evolutionary force such as genetic drift might act on a clonal population and contribute to generating LD in *Alternaria* spp. observed in this study.

## Conclusions

The GBS analysis of *Alternaria* spp. provided genome-wide SNP data and revealed genetic differentiation. The GBS method allowed effective comparison between *A. linariae* and *A. solani* and clearly distinguished them at a specific level by PCA and refined our understanding of the species responsible for the EB of tomato and potato. The *A. linariae* population from NC showed evidence of common ancestry and limited admixture and geographic isolation. Our data demonstrated non-random associations among SNP variants and these results were in line with the asexual reproduction of these pathogens. Furthermore, additional sampling may be necessary to determine host specificity and macroevolutionary patterns among these species in tomato and potato production regions. This study provides useful tools to plant pathologists and plant breeders in their efforts to develop resistant cultivars for commercial tomato and potato production in the United States.

## Supporting information

**Table S1. Isolates of *Alternaria* spp. collected from tomato and potato from North Carolina (NC) and Wisconsin (WI), respectively and used for genotyping by sequencing.** (XLSX)

## Acknowledgments

We thank the following people who provided technical assistance in this project: Inga Meadows, Virginie Rumsch and Stella Chang. We thank the editor and anonymous reviewers for their insightful and constructive comments for improving the manuscript.

## Author Contributions

Conceived and designed the experiments: TBA. Performed the experiment: TBA. Analyzed data: TBA BJK NJG. Contributed reagents/materials/analysis tools: TBA DH FJL BJK NJG. Wrote the paper: TBA. Reviewed and edited the paper: TBA BJK NJG DH FJL.

